# The conservation effectiveness of Indigenous and local community governance

**DOI:** 10.64898/2026.10.12.758350

**Authors:** Erika S. Zavaleta, Christine E. Wilkinson, Kelly Zilliacus, Blair C. McLaughlin, Bernard R. Tershy, Jahiya P. R. M. Clark, Roger Faust

**Author notes:** Research & Collections Department, Natural History Museum of Los Angeles County; Los Angeles, CA, 90007 USA and California Academy of Sciences, San Francisco, CA 94118 USA.

## Abstract

Effective governance of natural areas is central to conserving biodiversity and Earth’s life support systems, including climate regulation and food and water provisioning. Understanding the relative effectiveness of different governance options is thus critical. We conducted a global meta-analysis of 409 cases in 42 countries, comparing ecological outcomes over time in areas governed by Indigenous Peoples and local communities versus conventional governance approaches. Outcomes in community-governed areas matched those in conventional fully protected areas and exceeded those in partially protected and unprotected areas. Communities with legal land title had better outcomes than communities without title. We provide the strongest evidence to date that ecological outcomes of Indigenous and community governance meet or exceed those of other area-based conservation approaches and that titling strengthens these outcomes.

---

Efforts to conserve biodiversity increasingly depend not only on how much of the planet is conserved, but also on who governs those lands and waters. Accelerating threats to biodiversity, coupled with the argument that minimizing human pressures is the best way to protect it (*1*) have driven the global expansion of protected areas (*2*) controlled mainly by national governments (*3*). Meanwhile, Indigenous Peoples and local communities (“communities”) still manage and maintain rights to many of Earth’s most biodiverse areas, including over a quarter of global land surface and an even greater proportion of intact landscapes (*4*). The relative roles of these two distinct governance approaches in protecting biodiversity underlie a decades-long debate about how best to achieve conservation outcomes while sustaining human communities (*5*–*8*). Despite extensive research on both approaches, we lack a quantitative synthesis comparing conservation outcomes between community-governed areas and conventionally-governed protected areas (*9, 10*).

Comparing the conservation effectiveness of conventional and community governance has become especially important in light of international agreements that highlight the roles of each (*11*). Previous work examines the social and ecological effectiveness of either community-governed areas (*12*) or conventional protected areas (*13*–*16*). Others have qualitatively synthesized comparisons of ecological outcomes in Indigenous- and local community-governed areas versus protected areas, or have tallied which governance approach performed better across studies, without accounting for differences in effect sizes, study quality, and scope (*10, 12, 17, 18*). Existing quantitative, comparative syntheses have been limited to particular biomes and ecological outcomes (e.g. *19*). Finally, previous studies compare the conservation effects of different goals pursued by externally-governed projects, such as those targeting only biodiversity versus those with both social and ecological goals (*20*). However, goals differ from governance, which concerns who has authority and responsibility to set those goals and to make decisions about access, uses, management, and enforcement (*7, 21*).

To address how area-based conservation outcomes differ between community- and conventionally-governed areas, we conducted a global meta-analysis of published studies that directly compare the two approaches at country- or finer scales. From 39,182 retrieved papers, we retained studies that reported ecological outcome data over time (>1 year), for one or more cases comparing community- to conventionally-governed areas (fig. S1, see SI Materials and Methods for details). We categorized conventionally-governed areas as either fully protected, partially protected, or outside protection to enable policy-relevant comparisons (Fig. 1). In the community-governed group, we included areas governed or co-governed by both Indigenous Peoples and local communities, as both are recognized by international agreements (*11*) and much of the literature groups them together (*18*).

**Figure 1.**
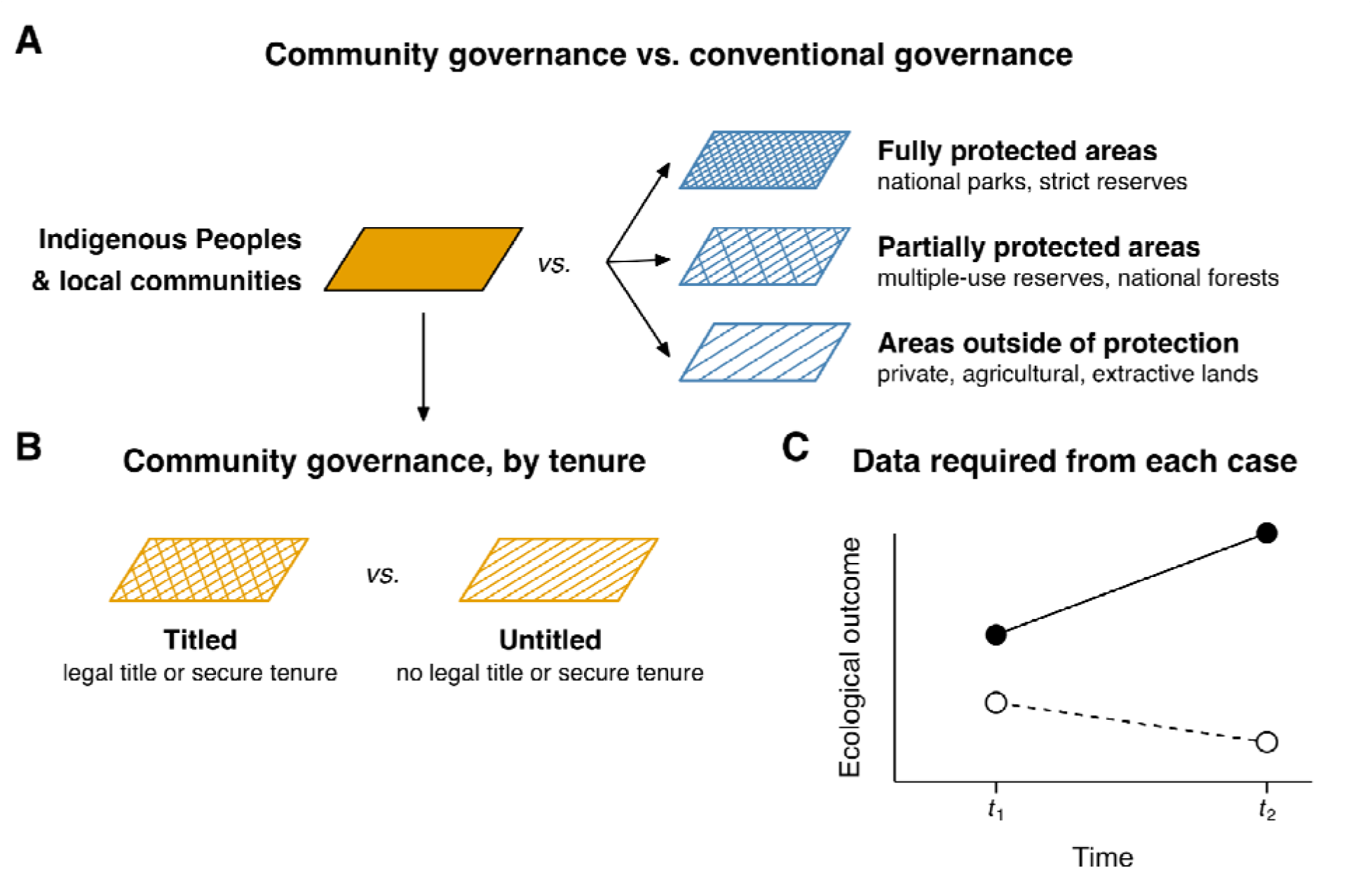
Design of meta-analyses comparing ecological outcomes between (A) areas governed by Indigenous Peoples and local communities (“community-governed”) and three classes of conventional area-based governance, with examples of each, and (B) community-governed areas with and without legal title or other forms of secure tenure. (C) Data extracted from each case included in our meta-analysis must have measured an ecological response variable at least twice, one or more years apart, in both community-governed (white points) and comparison (black points) areas with data reported at country- or finer-scales. Rates of change through time (slopes) are compared, in part so that pre-existing ecological differences between the community- and conventionally-governed areas are not misinterpreted as effects of governance type.

## Comparing community and protected-area outcomes

Across 409 cases in 42 countries (Fig. 2, fig. S2), ecological outcomes of community-governed areas were comparable to or stronger than comparison areas. Community-governed outcomes did not differ from those in conventional fully protected areas, such as national parks and strict biosphere reserves (Fig. 4, fig. S3, tables S1 and S2, p=0.43). Community-governed outcomes were better than outcomes in conventional partially protected areas, such as multiple-use reserves, game controlled areas and national forests (Fig. 4, fig. S4, tables S1 and S2, p=0.0035). Community-governed outcomes were also better than those in areas outside of conventional protection, such as privatized lands and state-governed areas designated for extraction, agriculture or other uses (Fig. 4, fig. S5, table S1 and S2, p<0.0001). For all three comparisons, restricting the set of cases to those employing matching techniques to further control for selection effects (*22*) generated no qualitative differences in findings (table S1, fig. S6) and were consistent with simple counts of cases in which each governance type outperformed the other (table S2).

**Fig. 2.**
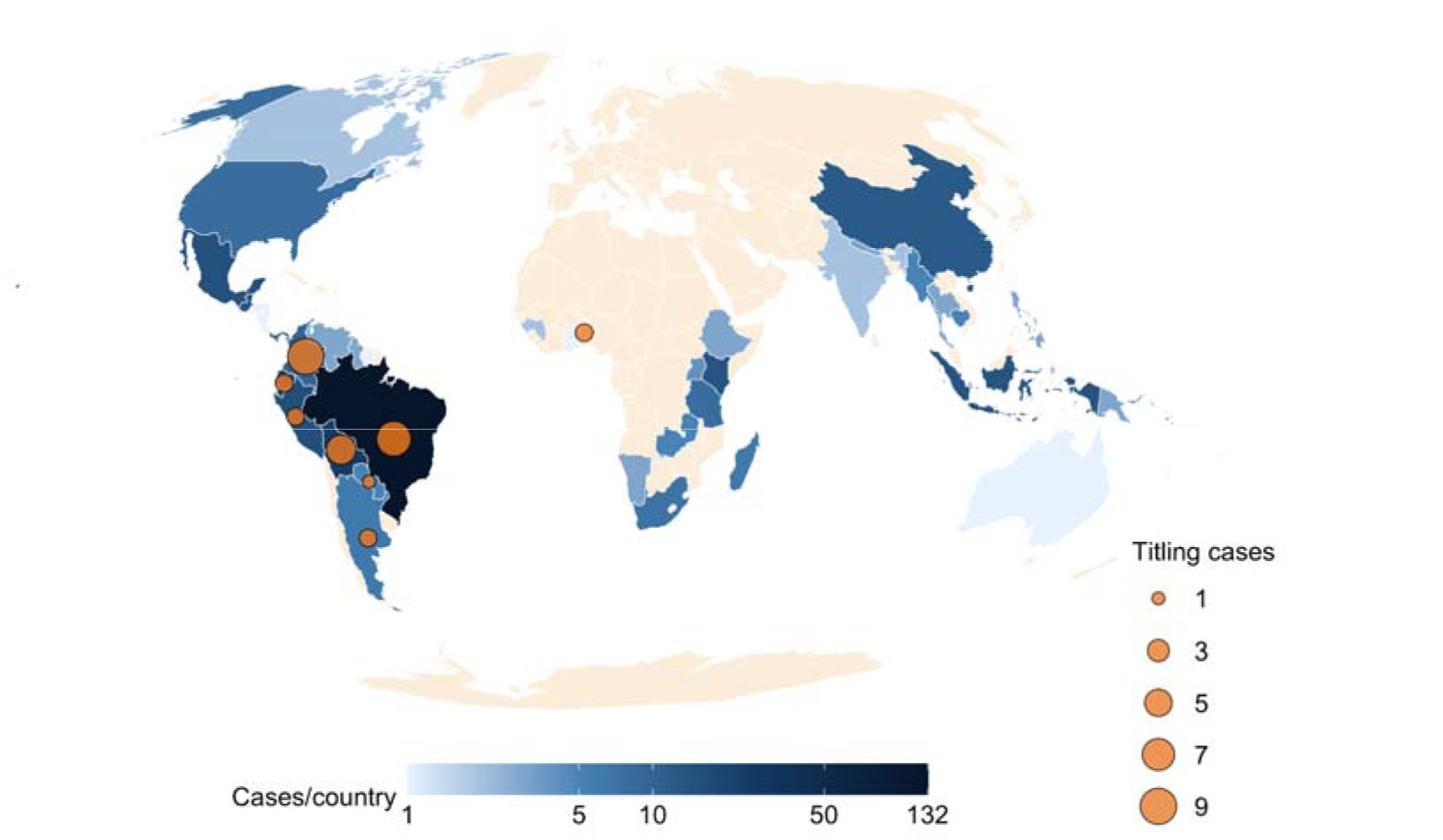
Global distribution of cases studying ecological outcomes in areas governed by Indigenous Peoples and local communities. (“communities”) and comparing them to conventional governance alternatives. Blue shading: cases comparing ecological change over time between community-governed areas and conventionally-governed fully protected areas, partially protected areas, or areas outside of protection. Orange circles: cases comparing ecological change over time between community-governed areas with and without formal land tenure or titles.

These patterns extended across most regions and ecosystem types (figs. S7 and S8). Most studies in our analysis measured habitat outcomes (Fig. 3A) and occurred in tropical forests (Fig. 3B). To maximize our power to detect influences of study attributes and additional ecological and social variables on effect sizes, we pooled our three conventional governance types (See Supplementary Text for details). No variable significantly explained variation in effect sizes (fig. S9), reflecting our sample structure’s low power (fig. S10). Nevertheless, community governance yielded better outcomes than conventionally governed comparison areas in all terrestrial ecosystems, while no significant differences between governance types were detected in marine systems (fig. S7). Similarly, community governance yielded better outcomes in South America, Africa and Asia, while no significant differences between governance types were detected in Oceania and North America (fig. S8). Community-governed outcomes were better than conventional governance for habitat-scale ecological variables, which are the type most cases measured, but not significantly better for species- and climate-specific variables (fig. S11).

**Fig. 3.**
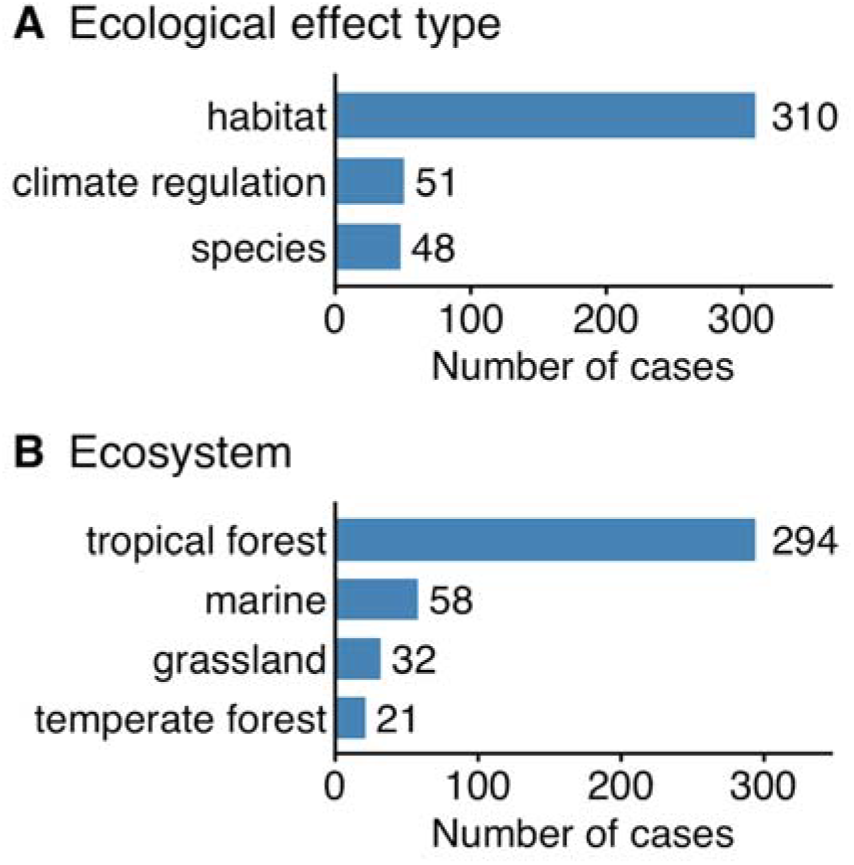
Cases measuring ecological effects of Indigenous and local community governance and comparison governance types, including fully protected, partially protected and unprotected areas by (A) type of ecological outcome measured and (B) ecosystem type studied. “Habitat” included variables such as forest and coral cover; “climate regulation” included variables such as avoided emissions and carbon stored in biomass; “species” included variables such as abundance, density or biomass of animal species, functional groups (e.g. predators) or taxonomic groups (e.g. carnivores).

## Outcomes of increasing tenure security

When we analyzed a second set of global cases that directly compare ecological outcomes in Indigenous and local communities with and without legal tenure or titles (Fig. 1, see SI text), titling of community-governed areas was associated with significantly better ecological outcomes than in areas without title (Fig. 2, Fig. 4, Table S1). Previous work corroborates this result, finding that environmental outcomes of securing tenure are positive in most cases in Central and South America (*23*) and across more of the globe from a small available sample (*24*). We found few quantitative, counterfactual (*25*) studies assessing the effects of securing title outside of South America, and all cases we were able to retain for this comparison addressed habitat outcomes in tropical forests.

**Fig. 4.**
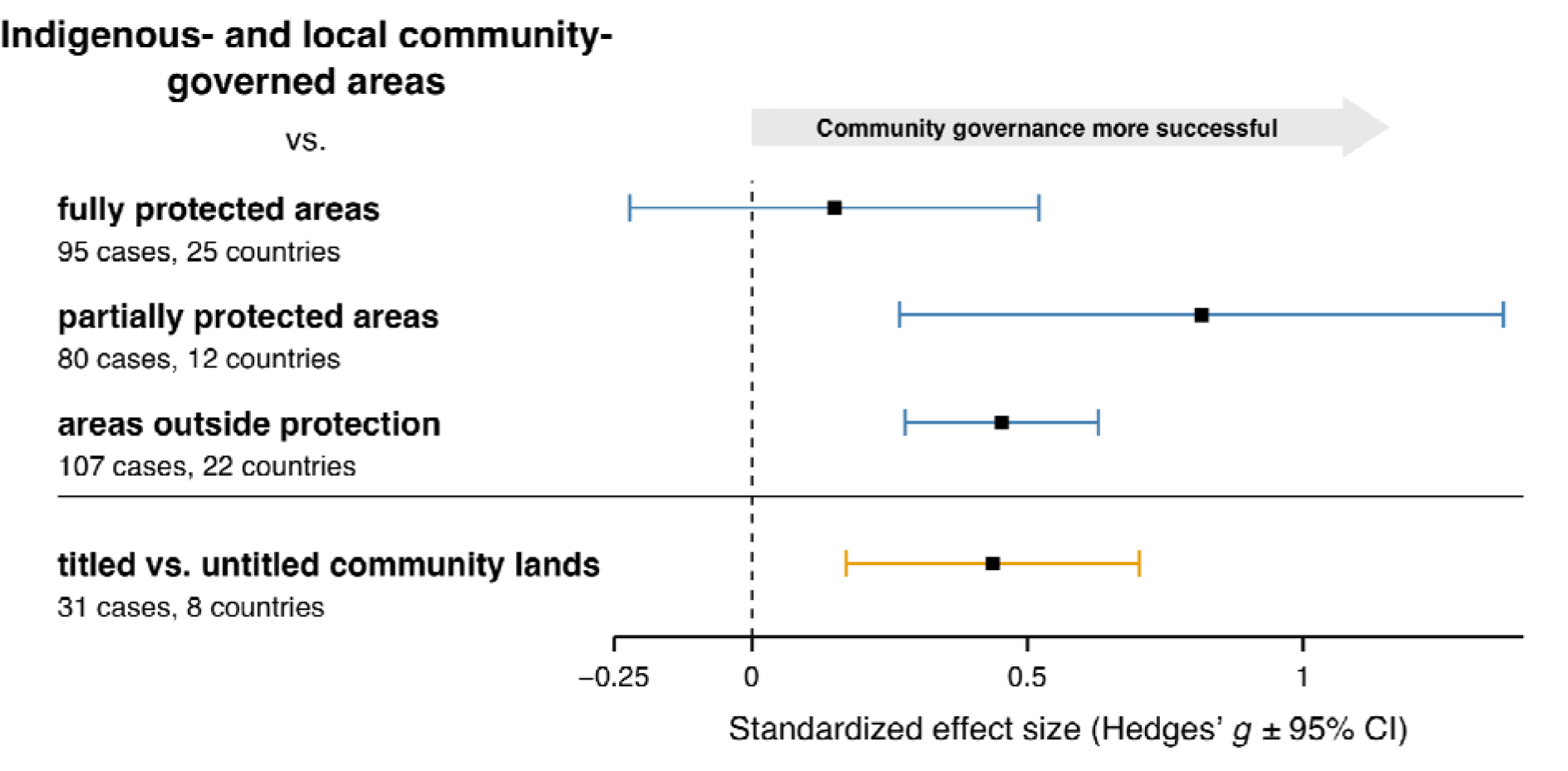
Effect sizes of Indigenous- and community-governed areas versus three conventionally-governed area types (top three bars), and effect sizes of titled (securely tenured) versus untitled Indigenous- and community-governed areas (bottom bar). Numbers of cases and countries in which cases occurred are reported for each comparison. Mean standardized effect sizes (Hedges’ g) are indicated by black squares, with 95% confidence intervals (CI) indicated by colored bars. The vertical dashed line indicates an effect size of zero. Values to the right of zero indicate that Indigenous- and community-governed area(s) had more positive ecological outcomes than the comparison group (blue bars), and titled community land had more positive ecological outcomes than untitled community lands (orange bar). Effect sizes are significant where the confidence intervals do not overlap the dashed zero line.

## Discussion

The positive ecological outcomes of conventional protected areas governed by state or private entities are extremely well documented (*13, 14, 16, 26*). However, we found they do not outperform areas governed by Indigenous and local communities when compared to them quantitatively across the global body of available counterfactual studies (*25, 27*). We provide the strongest evidence to date assessing the relative effectiveness of different forms of area-based conservation governance globally. Indigenous- and local-community governance performed as well as or better than other area-based conservation approaches. Strengthening Indigenous and community land tenure security further enhanced ecological outcomes.

Literature bias could influence the findings of our meta-analyses, arising from non-publication of studies lacking significant findings or author bias towards highlighting cases with a particular direction of effect. We found no evidence of significant publication bias in any direction in any of our four meta-analyses (table S3). However, consistent with previous work (*18*), cases comparing community-governed and fully protected areas reported fewer large, positive effect sizes for community governance than statistically expected (n=30 missing cases, table S3). This indicates that the estimated effect size of community governance is, if anything, conservative and increases confidence that results are not biased towards reporting of stronger outcomes for community-governed areas.

Consistent with the body of research on mechanisms, our findings reinforce others that titling can increase communities’ ability to enforce rules and legitimize them to outsiders (*23, 24*). A key determinant of all conservation governance outcomes is their effectiveness at rebuffing incursions by outside actors (*28*–*30*) and external, potentially mobile pressures including extractive industry and ecotourism development (*31, 32*). These can be strengthened considerably through national government policies, support through international agreements and mechanisms, and civil society. Efforts to bolster conventional protected area effectiveness are widespread and not yet matched with external support for community-governed areas, which in many cases protect their lands without compensation (*33*).

Our findings complement previous work on outcomes of different project goals (*20, 34*) and local participation (*35*) by addressing the effects of who has goal-setting and decision-making authority. Efforts aimed at local community benefits or participation without authority or title can have variable effects on their area-based governance and its effectiveness (*8, 36, 37*). For example, external imposition of rules on communities has been associated with weaker ecological outcomes and erosion of local institutions (*38*), suggesting that efforts to constrain or direct Indigenous and local community use and decision-making in their areas could be counterproductive. Conversely, our finding that legal title improves communities’ ecological outcomes indicates that bolstering these institutions can enhance conservation. The effects of titling depend on regional context and can erode community autonomy in some contexts (*39*). More counterfactual evidence is needed for titling’s effectiveness beyond South America (but see (*40*)) to better understand the place-based risks and benefits of strengthening tenure for conservation benefits.

Moreover, important geographic gaps exist in understanding community-governed conservation outcomes relative to other approaches. These include regions where Indigenous and local communities exercise extensive customary control, including in the world’s second largest tropical forested region in the Congo Basin, in Australia, and across most of the world’s northern high-latitude areas (*4*). Sze et al. (*19*) found that Indigenous lands experienced less deforestation than protected areas over time in Africa’s tropical forests and linked this in part to widespread, uncompensated community displacement for protected areas in the Congo Basin (*41*). However, more recent work calls this into question (*42*), underscoring the need for more clarity in this biologically and culturally diverse region. More broadly, the existing global body of evidence remains insufficient to robustly assess how study attributes, ecological characteristics, and socioeconomic factors influence effect sizes. Expanding counterfactual studies in underrepresented biomes, including marine systems, grasslands, and temperature forests; and incorporating underrepresented measures, particularly species- and biodiversity-level outcomes, could better inform action.

We focused on ecological outcomes of different conservation governance approaches; however, conservation success can be measured using a broader range of outcome variables (*43, 44*) that reflect a wider range of perspectives about what constitutes conservation effectiveness (*10*), including social as well as ecological outcomes (*45*). Positive social outcomes have been documented extensively in association with Indigenous- and community-led stewardship (e.g. (*12, 18, 46*) and strengthening of community tenure security (*24*). Social benefits are also positively associated with desired ecological outcomes in a global protected-area study (*15*), indicating that perceived tradeoffs or conflict between ecological and social conservation outcomes might be exceptions rather than the norm.

At present, the vast majority of formally recognized protected areas worldwide are controlled by governments, with only a small fraction governed by Indigenous Peoples and local communities (*3*). Many national governments recognize conventional protected areas, but not Indigenous- and community-governed areas, as contributing to current international conservation targets including 30x30 (*47*). Many national governments and aid organizations also direct substantially more funding toward conventional protected areas than for Indigenous- and community-governed areas (*33*), leaving local communities to bear the labor and other costs of conservation (*48*).

Our findings have important implications for how conservation is recognized, supported, and financed. Our finding that Indigenous- and local community-governed areas had equal or greater conservation outcomes than conventional protected areas strongly supports counting these Indigenous and community areas towards international conservation targets. Our finding that titling strengthened conservation outcomes supports the use of legal community tenure as a conservation tool. Finally, because community-governed areas deliver comparable or better outcomes despite receiving less support, bringing government and aid investment in line with their conservation contribution could yield greater conservation gains.

## Acknowledgments

We thank Morgan Abbott, Kyle Bobiwash, Solange Bandiaky-Badji, João Campos-Silva, Teo Cooper, Nicolàs Cruz, Valeria Cuevas, James Deutsch, Alain Frechette, Reza Goljani Amirkhiz, Graden Froese, Nautica Jones, Niko Kaplanis, Kyaw Kyaw, Julianna Rossi, Nonette Royo, and Peter Seligmann for valuable comments.

## Author contributions

Conceptualization: ESZ, CW, BRT

Methodology: CW, ESZ

Investigation: ESZ, CW, KZ, JC, BCM

Visualization: ESZ

Funding acquisition: ESZ

Project administration: ESZ, CW, KZ

Supervision: ESZ

Writing – original draft: ESZ, CW, BCM, KZ, JC, RF

Writing – review & editing: ESZ, CW, BRT, JC, KZ, BCM, RF

## Data, code, and materials availability

All data and code will be available on Dryad at the time of manuscript publication, and the citation and link added here and in the references.

## Supplementary Materials

Materials and Methods

Supplementary Text

Figs. S1 to S11

Tables S1 to S3

References 50-64

