## Supplementary Materials for "The conservation effectiveness of Indigenous and local community governance"

**The PDF file includes:**

Materials and Methods

Supplementary Text

Figs. S1 to S11

Tables S1 to S3

References

**Materials and Methods**

We preregistered our hypotheses, sampling plan, and analysis approach  on the Open Science Framework (OSF) on July 9, 2025 ([https://doi.org/10.17605/OSF.IO](https://doi.org/10.17605/OSF.IO/XXXXX)/CJD3S) prior to screening papers and beginning analysis. We modified our preregistered approach in the following ways: we performed additional publication bias tests, sensitivity analyses and simulations as described below. We refrained from imputing ecological outcomes from cases missing measures of central tendency (e.g. means, medians) for ecological outcomes and instead excluded studies that did not report means or other measures of central tendency. We refrained from using AIC to compare models fitted using different approaches (e.g. ML vs. REML).

***Literature Search***

We conducted a systematic review of the English-language literature using two databases: Web of Science and ProQuest. We conducted the search using three four-keywords sets, with sets grouped into 1) Indigenous or local community steward keywords, 2) management keywords, 3) contextual keywords, and 4) impact keywords, with each group separated by an 'AND' operator.

Indigenous or local community stewardship keywords were: ("Indigenous" OR "local community" OR "community-based" OR "community based" OR "traditional ecological" OR "ancestral" OR "First Nations" OR "locally based" OR "locally-based" OR "rights-based" OR "rights based" OR "decentraliz*" OR "decentralis*" OR "community forest*" OR "communal land" OR "local stewardship" OR "community fishery management" OR "community fisheries management")

Management keywords were: ("management" OR "governance" OR "self-governance" OR "self-governing" OR "stewardship" OR "tenure" OR "monitoring" OR "custodian*" OR "custody" OR "guardian*" OR "fishery" OR "fisheries" OR "forestry" OR "conservation" OR "protection" OR "protected" OR "practice*" OR "co-management" OR "collaborative management" OR "reserve" OR "reserves" OR "conservanc*" OR "restoration" OR "reintroduction" OR "ecotourism" OR "eco-tourism" OR "wildlife tourism")

Contextual keywords were: ("wildlife" OR "species" OR "habitat*" OR "forest*" OR "wetland" OR "fish*" OR "nature" OR "natural" OR "river" OR "lake" OR "stream" OR "water" OR "ocean" OR "tree*" OR "ecosystem" OR "ecolog*" OR "rangeland*" OR "land cover" OR "land*" OR "deforest*" OR "mammal*" OR "plant*" OR "animal*" OR "freshwater" OR "overharvest*" OR "biodiversity" OR "carbon" OR "multispecies" OR "multi-species" OR "environment*" OR "phytoplankton" OR "plankton" OR "biomass" OR "grassland*" OR "savanna" OR "rainforest*" OR "mountain*" OR "natural resource*")

Impact keywords were: ("impact*" OR "effective*" OR "benefit*" OR "co-benefit*" OR "co-benefits*" OR "reduction" OR "recover*" OR "resilience" OR "improve*" OR "increase*" OR "reduc*" OR  "sustain*" OR "positive outcome*" OR "growth" OR "cost-effective" OR "regenerat*" OR "protect*" OR "conserv*" OR "inhibit*" OR "positive change*")

In Web of Science, we retained results within the following categories: environmental sciences, environmental studies, ecology, biodiversity conservation, green sustainable science tech, water resources, plant sciences, forestry, geography, multidisciplinary sciences, marine freshwater biology, agronomy, social sciences interdisciplinary, regional urban planning, zoology, fisheries, evolutionary biology, urban studies, economics, geosciences multidisciplinary, biology, microbiology, oceanography, sociology, public environmental occupational health, entomology, anthropology, soil science, geography physical, development studies, international relations, infectious diseases, agriculture multidisciplinary, horticulture, toxicology, ethnic studies, remote sensing, limnology, political science, area studies, humanities multidisciplinary, ethics, geology, mycology, agriculture economics policy, cultural studies, archaeology, women's studies, language linguistics, linguistics, religion.

In ProQuest, we retained results within the following categories: biodiversity, conservation, climate change, Indigenous peoples, community, sustainability, ecosystems, sustainable development, forests, natural resources, economics, invasive species, rivers, studies, resource management, Native peoples, decision making, plant species, fish, environmental impact, land use, wildlife conservation, rural areas, community involvement, case studies,  agriculture, freshwater, environmental protection, ecology, fisheries, vegetation, forestry, habitats, forest management, native species, nonnative species, households, marine, communities, decentralization, flowers and plants, trees, local communities, population, urban areas, tourism, food, management, knowledge, species diversity, governance, native organisms, environmental policy, water quality, culture, Indigenous knowledge, economic development, collaboration, species richness, farmers, seeds, developing countries--ldcs, fishing, environmental restoration, politics, water management, research, data collection, restoration, biomass, microorganisms, policies, protected areas, anthropogenic factors, deforestation, soils, lakes, adaptation, intervention, carbon, national parks, genetic diversity, [economic development: agriculture, natural resources, energy, environment, other primary products (o13)], Indigenous populations, livestock, nature conservation, environmental management, wildlife, islands, abundance, poverty, local government.

These searches returned 55,029 records, yielding 39,812 records after duplicates were removed.

***Study screening process***

We used Covidence systematic review software (www.covidence.org) to screen titles and abstracts. We followed the PRISMA guidelines for conducting a meta-analysis, and followed the SAFE procedure for stopping criteria when screening abstracts (*50, 51*). We also used the Covidence relevance ranking algorithm to determine an initial cut-off point for gathering relevant literature. Specifically, in Phase 1 we screened a random set of training data comprising at least 1% of the total number of records. In Phase 2 we applied active learning in batches of 100 records until the following criteria were met: 1) at least twice the number of training records were screened, 2) a minimum of 10% of the total dataset was screened, and 3) we screened at least 5,000 records using the relevance ranking algorithm. In Phase 3, we conducted forward-backward citation searches on relevant reviews and meta-analyses identified during the abstract screening process. All 402 records resulting from this screening process were reviewed in full.

*Study inclusion criteria*

1. Indigenous Peoples and local community governed (hereafter community-governed) areas assessment: studies needed to assess stewardship of areas governed by Indigenous Peoples or local communities. We relied primarily on authors’ characterization of Indigenous and/or local community-governed. We excluded cases that identified communities as settlers, recently relocated groups, or those in which land was privately owned by individual households. The authors of studies included in this meta-analysis often did not specify whether or how they identified the communities whose lands they studied as Indigenous or non-Indigenous (see also *18*). We thus did not attempt to distinguish Indigenous-led from local community-led governance.

2. Study design: studies needed to compare at least one community-governed area to at least one fully protected, partially protected, or unprotected area; or compare titled and untitled community-governed areas. We relied primarily on authors’ characterization of fully protected areas, partially protected areas, and unprotected areas, and titled versus untitled community-governed areas.  Protected areas included those operated by states as well as non-governmental or private organizations or concessionaires. Comparisons of titled and untitled community-governed lands were identified by study authors as studies of the effects of titling; secure, formal or legal land tenure; homologation; or legal dedication.

 The following study types were included:

A. Experimental: studies that had random assignment to groups with replication. (None of this type were identified in our screen.)
A. Quasi-experimental: Studies that lacked random assignment to intervention and counterfactual groups but that accounted for differences between them to create comparison groups with similar attributes.
A. Intervention vs. counterfactual: Studies that compared the value of an ecological variable in community-governed area(s) (intervention) to conventionally-governed or untitled comparison area(s) (counterfactual) over a particular time period, without explicitly estimating differences in before-and-after changes between groups.
A. Before-after-control-intervention: Studies that used both before vs. after and control vs. intervention in their design but did not use statistical matching.
A. Before-after studies for which there was both an intervention and a counterfactual in the same period, over time, accounting for differences in when areas received the intervention (i.e., (*52*) - difference-in-difference analysis of staggered receipt of land tenure).

3. Response variables: studies needed to measure at least one ecological effect. When two or more response variables in the same study were judged on theoretical grounds to be highly correlated (positively or negatively), we selected the most ecologically relevant one.

4. Time period: studies needed to measure at least one ecological effect at at least two time points, reported as change over time, start and end values, or overall difference between rates of change of the community-governed areas and the comparison area type.

5. Data reporting: studies needed to report data with means and central tendencies. If a study did not include measures of central tendency for ecological effects, such as means or medians, we departed from our registered protocol by excluding them rather than imputing these values. If standard deviations were missing, we attempted to estimate them using alternative measures of dispersion (standard errors, confidence intervals, t-values) if possible.

We excluded multi-country or global analyses in which data were not provided separately by countries; meta-analyses and literature reviews; and studies with unspecified start and end years that preclude determination of the study duration, unless a study provided duration and/or an annualized measure of change (e.g. in a differences-in-differences analysis examining 3 years after designation) along with a range of dates within which all analyses fell.

***Data extraction and quality control***

*Data screening and extraction*

Data screening outcomes are summarized in figure S1 (*53*). Four authors contributed to screening titles and abstracts, and five authors contributed to screening the full text of papers retained in the title/abstract stage. At the full-text stage, all papers were read by two authors to determine eligibility, with disagreements settled by a third author (CW or ESZ) and then by a meeting to discuss remaining edge cases.

*Case designation*

When studies measured ecological change via more than one metric, we treated each as a distinct case nested within the study. For studies that had multiple conventional governance types, we treated each community/conventional governance combination as a separate case nested within study. For studies with analyses covering multiple countries and with data reported by country, we treated cases from each country as separate and nested within study.

*Data extraction*

For each case we extracted the following information: the ecological effects measured, which we then categorized into 3 ecological effect types: habitat, climate regulation, and species; ecosystem type; geographic region; study start and end years; comparison area type (fully protected, partially protected, unprotected); sample size; and autonomy score for the community-governed area(s) (*54*).  Each case was scored for autonomy independently by two reviewers, and when the two reviewers disagreed, a third reviewer assessed which score to assign. Data extraction was repeated by a second author for twenty randomly selected papers to check for accuracy. Finally, two authors (ESZ, KZ) reviewed all retained cases for completeness of extracted data and conformance with inclusion criteria.

All ecological effects measured were evaluated by screeners as positive (an increase in the measure is desirable, e.g. wildlife density) or negative (an increase in the measure is not desirable, e.g. deforestation rate). Outcome values for negative ecological effects were multiplied by -1 to yield consistency in the correspondence of positive values with desirable or positive ecological outcomes. Two cases were omitted from the study because the response variable could not be confirmed as positive or negative by the reviewer team.

*Sample size calculation*

To determine sample size (n) for each case, we included replicates that estimated central tendency and dispersion of an overall ecological response. For example, if a study’s reported sampling unit was a pixel, and each pixel was assigned a single landcover value, we did not consider n to equal the number of pixels even if that was reported as n. If a study’s sampling unit was a plot within a reserve, within which a central tendency and measure of dispersion of a rate, density or other dependent variable was estimated quantitatively, we considered it a replicate contributing to n. When sample sizes were not provided, we first attempted to count them from a study’s maps and figures, and then if possible searched for n (e.g. in a study of “all Indigenous Territories” in a particular country or state) in authoritative sources or databases for the described area type and geography. We used PlotDigitizer for extraction of sample sizes and data from figures, and Fiji (Image-J) when necessary to accurately count study sites depicted as polygons or points on maps.

*Additional data*

For each case, we assigned the following country-level indices taken in the end year of each study: wealth (log GDP, Penn World Table v. 11.0) (*55*); inequality (SWIID 9.92 Gini index) (*56*); WGI Control of Corruption and Rule of Law estimates (*57*). Control of Corruption measures things like whether bribes are taken, while Rule of Law measures more whether laws (including environmental laws) are enforced. We extracted absolute governance scores (range 0-100, a score’s meaning is unchanging across years to allow comparison across years) of each for the end year of each study (since, in part, the indices go back to 1996 and many studies began before that). Studies that ended in 1999 were assigned a value from 1998 because of missing data in the WGI dataset. Low scores for both metrics are indicative of lower governance functioning.

***Analyses***

All analyses were conducted in R using the RStudio interface (v. 2026.09.0+174). We used artificial intelligence (Claude, Opus 5) interactively to support code writing, debugging, revision and formatting, and we checked and edited all code manually by line and section.

*Effect size calculation*

We measured ecological outcomes as effect sizes (Hedges’ g). Hedges’ g, a standardized mean difference, was computed as *((mean rate of change for intervention area – mean rate of change for control)/pooled standard deviation)*bias correction factor for small sample sizes,* where the intervention is the community-governed or titled community value, and the control area is the conventionally governed, or untitled community value, in each meta-analysis type we report, respectively. The pooled standard deviation combines the standard deviations of the two groups (community-governed areas and comparison governance or titling type) for the ecological metric in each study. Hedges’ g effect-size equations (Eqs. 1-3) are:

$$g=J\cdot\frac{\bar{X}_{T}-\bar{X}_{C}}{S_{\text{pooled}}}$$

$$S_{\text{pooled}}=\sqrt{\frac{\left( n_{T}-1 \right)s_{T}^{2}+\left( n_{C}-1 \right)s_{C}^{2}}{n_{T}+n_{C}-2}}$$

$$J=1-\frac{3}{4\left( n_{T}+n_{C}-2 \right)-1}$$

where J is the small-sample bias correction, X values are means per group, S is standard deviation, s^2^ is variance per group and n is sample size per group. The variance of Hedges’ g is (Eq. 4):

$$v_{g} =\left( n_{I}+n_{C} \right)/\left( n_{I}*n_{C} \right)+\left( g^{2} \right)/\left( 2*\left( n_{I}+n_{C} \right) \right)$$

Hedges’ g has several advantages as a meta-analytic metric, including that it is well equipped to deal with variation among cases in distribution structure and variance through use of their standard deviations. For cases where standard deviations were not reported, we extracted and converted to standard deviation values other dispersion measures including standard errors, confidence intervals and root mean squared errors. Although we searched for p-values and other test statistics such as t and Z where no dispersion measures were reported, we identified no cases where they were suitable for use. For cases lacking any measure of dispersion, we imputed pooled standard deviations as the median of available values. We stratified available values and recipient cases into two data-type groups, proportional (e.g. percentages) and non-proportional (e.g. hectares, counts) to reflect the distinct dispersion structures of these different response variable types. Our sensitivity checks included model runs excluding cases with imputed dispersion values. Because Hedges’ g has a variation that mechanically varies positively with g^2^, recommended tests for publication bias correct

Hedges’ g also allows inclusion of data from comparisons of effects over time reported in multiple ways. We classified each case into one of three data reporting types. Type 1 studies reported full Before-After-Control-Impact data, including separate data values for the start and end timepoints for each of the community-governed areas and the comparison areas (fully protected areas, partially protected areas, areas outside protection). Type 2 studies reported rates for each treatment without providing separate start and end values.Type 3 studies reported the difference in rates between the community-governed and comparison areas without providing separate rates for each. Because Hedges’ g cannot be calculated for cases with n=1 sample per treatment or with missing n, we ran sensitivity checks and comparison models using the effect size ln(RR), which allows inclusion of n=1 and missing-n cases but excludes cases with Type 3 data (see *Sensitivity Checks* below).

*Vote-count classification*

While vote-counting approaches obscure variation in effect sizes, dispersion, study scope and quality, we include them as a qualitative check on the meta-analytic effect-size results. For each comparison (community-governed areas vs. conventional fully protected areas, conventional partially protected areas, or areas outside of conventional protection; titled vs. untitled community areas), we tallied the number of cases in which community-governed area(s) had a more positive effect than the comparison area(s), and the number of cases in which the comparison area(s) had a more positive effect than the community-governed area(s).

*Main meta-analytic models*

We used multi-level random-effects meta-analytic models to test the hypothesis that ecological outcomes under community-governed areas differ from those under alternative governance types. We ran separate models of studies comparing outcomes over time from community-governed areas with those of 1) fully protected areas, 2) partially protected areas, and 3) areas outside of conventional protection. We also tested the hypothesis that ecological outcomes differ between titled and untitled community-governed areas, by running the models on studies that directly compared titled and untitled community-governed areas. We included study as a random factor in our models to nest the structure to reflect higher associations among cases within the same study.

To test whether each mean effect size differed from zero, we used a z-test with a two-tailed significance level of alpha=0.05. We quantified heterogeneity across studies using 1) Cochran’s Q-test to test for significant variation among effect sizes and 2) the multi-level I^2^ statistic to estimate the proportion of total variance attributable to within- and between-study heterogeneity (*57*). We report 95% confidence intervals around each effect size, considering results statistically significant if the confidence interval does not include zero. For studies with pre- and post- values both reported (Type 1 and Type 2), we use pooled (community-governed and comparison areas) pre-test standard deviation (*58*) and assume pre-post standard deviation correlation of rho=0.5 with sensitivity checks varying that assumption.

*Sensitivity checks*

We set a lower sample-size bound of 15 cases for sensitivity checks. We ran the following sensitivity checks for each main comparison group and the comparison of titled and untitled community lands:

Tests for publication bias: We performed Egger’s regression test with modified standard error (SE), the recommended approach to address the correlation of Hedges’ g with its own SE inherent in the way its calculated variance (SE^2^) (Eq. 4) (*58*). We applied multilevel PET and PEESE tests to account for the structure of cases clustered within studies (*59*).We used Duval and Tweedie’s (*61*) trim-and-fill tests to estimate numbers of missing papers for each meta-analytic comparison. If missing cases were identified, we refitted a basic random-effects meta-analytic model with the imputed missing cases included.

Leave-one-out checks: We checked the influence of individual studies on grand mean effect size of leaving out one study at a time (LOSO), and the influence of individual countries on grand mean effect size and 95% confidence intervals of leaving out one country at a time (LOCO).

Subsetting: To check for the effect of imputing missing dispersion values, we analyzed the subset of cases that reported dispersion statistics. Because studies employing matching procedures produce better estimates of intervention effects by controlling for effects of variables such as terrain, climate and proximity to population centers (*63*), we analyzed the subset of cases that used matching techniques, such as propensity-score matching and covariate matching. Because a large proportion of cases come from the country of Brazil, we analyzed the subset of cases not located in Brazil.

BACI assumptions about correlation between pre- and post-timepoint measures of dispersion: For Before-After-Control-Impact designs (Type 1), we re-ran two sensitivity checks, with the assumed correlation between pre- and post-dispersion values adjusted to r=0 and r=0.8 (to compare to the main model assumption of r=0.5), respectively.

lnRR analysis of cases excluded from calculation of Hedges’ g: We conducted analysis of the meta-analytic response metric ln(RR), which permits Type 1 and Type 2 cases with sample size n=1 or missing n values to be included, though it lacks the advantages of Hedges’ g for a study like ours in which response metrics vary, some response measures equal or approach zero, and sample sizes are highly variable (6*4*). We analyzed (1) only the set of cases that had been excluded from Hedges’ *g* computation, and (2) all cases for which ln(RR) could be computed (all except Type 3 cases reporting only an overall effect size, and cases with a zero value for the comparison group in the denominator).

*Meta-regression on moderator variables*

For community-governed  vs. conventionally-governed area comparisons, we analyzed the influence of the following potential moderator variables of effect size: comparison area type, study duration, study end year, geographic region, ecosystem type, ecological effect type, autonomy of governance (*54*), and the following country-level country-level indices taken in the end year of each study: log GDP (wealth), Gini index (inequality),WGI Control of Corruption and Rule of Law estimates.

To maximize power to detect effects of these potentially moderator variables, we pooled cases across the three comparison groups (community-governed areas versus fully protected, partially protected and unprotected) for the meta-regression. To assess our ability to detect effects of these moderator variables given our sample size, range of effect sizes and variances, and covariance matrix, we built power curves through simulated model runs (n=2000) across a range of hypothetical moderator effects. For each moderator variable we report the minimum effect detectable with 80% power.

We modeled effect sizes pooled across all three comparison groups (fully protected, partially protected, and unprotected) on the suite of moderator variables, retaining only one of any pair of variables with correlation >0.7 based on theoretical grounds and inspection of bivariate fits (rule of law vs. control of corruption was the only such decision). We applied cluster-robust analysis to account for clustering of cases within studies, and assumed a correlation of rho=0.5 between effect sizes arising from the same study (with sensitivity checks using rho=0.2 and 0.8). We tested for overall differences between subgroups using Q-tests for heterogeneity. If significant (p<0.05), we conducted pairwise comparisons between subgroups, applying Bonferroni corrections to adjust for multiple comparisons and reduce the risk of Type I errors.

For the comparison between titled and untitled community-governed areas, we were limited by the small size of the dataset to analysing a maximum of 3-4 moderator variables. We dropped the ecosystem type variable because all cases were in tropical forest, the region variable because nearly all cases were in South or Central America, and the autonomy variable because nearly all cases were scored at one of the top two autonomy levels, providing little variation. We began with duration and country-level indicators (control of corruption, rule of law, inequality and wealth), dropping the weaker bivariate fit of any pair of variables correlated at  > 0.7.

**Supplementary Text**

***Tests for publication bias***

Tests for publication bias were statistically insignificant (p>0.05) for each of the four meta-analyses (table S3). The community-governed areas vs. fully protected areas- analysis estimated 28 studies missing on the *right* side of the funnel plot, signifying potential bias against studies with large positive effect sizes. Estimation of weighted mean effect size with the inferred missing studies imputed yielded an overall g of 0.539 (random effects model, 95%CI 0.2820 to 0.7959, p<0.001), suggesting publication bias might have led to an underestimate of positive community-governed area outcomes relative to fully protected areas. The community-governed areas vs. areas outside protection analysis estimated 15 studies missing on the *left* side of the funnel plot, signifying that studies with small or negative effect size might be missing from the literature (potential publication bias). However, estimation of weighted mean effect size with the inferred missing studies imputed yielded an overall g of 0.218 (random effects model, 95%CI 0.0547 to 0.381, p=0.0089), indicating that our finding is robust to publication bias.

***Sensitivity checks***

For the community-governed areas vs. fully protected areas comparison, LOSO caused grand mean *g* (0.150, 95% CI  -0.221 to 0.521, n=95) to vary between 0.057-0.23, and LOCO caused it to vary between 0.047-0.40, with 95% CI overlapping 0 in every check. Exclusion of cases with imputed dispersion values and variation of the assumed correlation between pre- and post- dispersion values for BACI studies did not qualitatively affect results. Exclusion of cases in Brazil and restriction to matched cases did not qualitatively affect results; however, exclusion of Brazil increased the grand mean effect size g (Brazil-excluded g = 0.401 (95% CI: -0.150 to 0.951, n=63; matched cases only: g=0.104, 95% CI: -0.308 to 0.517, n=36, fig S6). Analysis using the effect size ln(RR) of available cases excluded from the Hedges’ g analysis (n=19) and of all eligible cases (n=102) did not change the direction or absence of significance of the result.

For the community-governed areas vs. partially protected areas comparison, LOSO caused grand mean *g* (0.816, 95% CI 0.268 to 1.36, n=80) to vary between 0.68 to 0.93, and LOCO caused it to vary between 0.58-0.92 with 95% CI overlapping 0 in each check except exclusion of Brazil (see below). Exclusion of cases with imputed dispersion values (n=57 cases excluded) did not qualitatively affect the estimate of grand mean effect size but widened the 95% CI to include 0, as did variation of the assumed correlation between pre- and post- dispersion values for BACI studies and exclusion of Brazil. Restriction to matched cases increased the grand mean effect size (g=1.165, 95% CI: 0.265 to 2.065, n=40, fig S6). Only 14 studies were omitted from the main analysis that were eligible for ln(RR) meta-analysis. Analysis using the effect size ln(RR) of all eligible cases (n=74) increased the 95% CI lower bound slightly to overlap zero but did not change the direction of effect (p=0.069).

For the comparison of community-governed areas vs. conventionally governed areas outside protection, LOSO caused grand mean *g* (0.453, 95% CI 0.278-0.628, n=107) to vary between 0.40 to 0.49, and LOCO caused it to vary between 0.40-0.48 with 95% CI overlapping 0 in every check. Exclusion of cases with imputed dispersion values (n=51 cases excluded) increased the estimate of grand mean effect size but widened the 95% CI to include 0. Neither variation of the assumed correlation between pre- and post- dispersion values for BACI studies nor exclusion of Brazil qualitatively affected the result. Restriction to matched cases increased the grand mean effect size (g=0.954, 95% CI: 0.072 to 1.836, n=42, fig S6). Only 14 studies were omitted from the main analysis that were eligible for ln(RR) meta-analysis. Analysis using the effect size ln(RR) of available cases excluded from the Hedges’ g analysis (n=50, no change in direction, p=) and of all eligible cases (n=132) did not qualitatively affect the direction or significance of the result.

For the comparison of titled and untitled community-governed lands, LOSO caused grand mean *g* (0.437, 95% CI 0.171 to 0.703, n=31) to vary between 0.36 and 0.41, and LOCO caused it to vary between 0.31-0.51 with 95% CI overlapping 0 in every check. Exclusion of cases with imputed dispersion values left only 12 cases, and only one study required an assumption about BACI dispersion, so these tests were omitted. Only 15 matched cases were available, and analysis restricted to them did not change the direction of effect but caused the 95% CI to overlap 0 (g=0.271, 95% CI: -0.131 to 0.674, n=15). Only 1 study was omitted from the main analysis that was eligible for ln(RR) meta-analysis, so this test was skipped. Analysis using the effect size ln(RR) of all eligible cases (n=21) did not affect the direction or significance of the result.

***Heterogeneity***

All meta-analyses had high heterogeneity, reflecting substantial expected variation in effect sizes across widely varying contexts. For the comparison of community-governed areas vs. fully protected areas, heterogeneity Q(94) = 8192 (p < 0.001; I²total = 99.0%). Between-study variance was σ² = 0.82 (95% CI 0.36 to 1.68) and within-study variance was σ² = 0.65 (95% CI 0.44 to 0.99). The 95% prediction interval was -2.26 to 2.56. For the comparison of community-governed areas vs. partially protected areas, heterogeneity Q(79) = 3560 (p < 0.001; I²total = 99.5%). Between-study variance was σ² = 1.11 (95% CI 0.26 to 2.75) and within-study variance was σ² = 2.62 (95% CI 1.79 to 3.98). Most of the heterogeneity (69.9%) occurred among effect sizes within studies. The overall effect was positive (0.82, 95% CI 0.27 to 1.36; Satterthwaite p = 0.007), and the 95% prediction interval (-3.01 to 4.64) indicates that true effects in different contexts may be negative or positive, as expected. Finally, heterogeneity Q(106) = 3648 (p < 0.001; I²total = 97.7%) for the comparison between community-governed areas and areas outside conventional protection. Between-study variance was σ² = 0.097 (95% CI 0.005 to 0.345), while within-study variance was σ² = 0.333 (95% CI 0.224 to 0.506). Once again, most heterogeneity (75.6%) occurred among effect sizes within studies, and the 95% prediction interval (-0.84 to 1.75) indicates that true effects in specific contexts may be negative or positive, as expected.

***Pooled moderator analysis***

No variable included in the pooled moderator model was significantly related to effect size at the p<0.05 level, though power analysis by simulation indicates this finding should be interpreted as absence of evidence rather than affirmative evidence that these moderators lack influence on the relative outcomes of community and conventional governance areas. Four contrasts had trends at the p<0.10 level: community-governed areas tended to outperform unprotected and partially protected areas more strongly than they outperformed fully protected areas; and community-governed areas tended to perform more strongly relative to comparison areas in temperate forests and temperate grasslands than in marine ecosystems (fig.  S9, n=218 effect sizes, 68 studies, 26 countries; cases missing more than 25% of moderator variable values omitted from analysis).

We lacked power to detect effects of most moderators; only three moderators (study duration, Gini coefficient and autonomy score) achieved 80% power to detect an effect of 0.5 in Hedges’ *g* given a 1 SD change in the moderator value (fig. S10). Two more moderators (control of corruption, final year of study) achieved 80% power to detect an effect of 0.8 in Hedges’ *g*. With grand mean effect sizes in the range of 0.2-0.8 in the main meta-analyses for each comparison group, our power to detect the effects of any moderator was small even for these six variables. Three key moderator effects - region, ecosystem type and ecological outcome type measured (e.g. habitat vs. species measures) - had reasonable power to detect only differences much larger than the overall grand mean effect sizes we found across cases, possibly in part due to the dominance of tropical forests, habitat-scale response variables and South America cases in the dataset.

**Supplementary Figures**

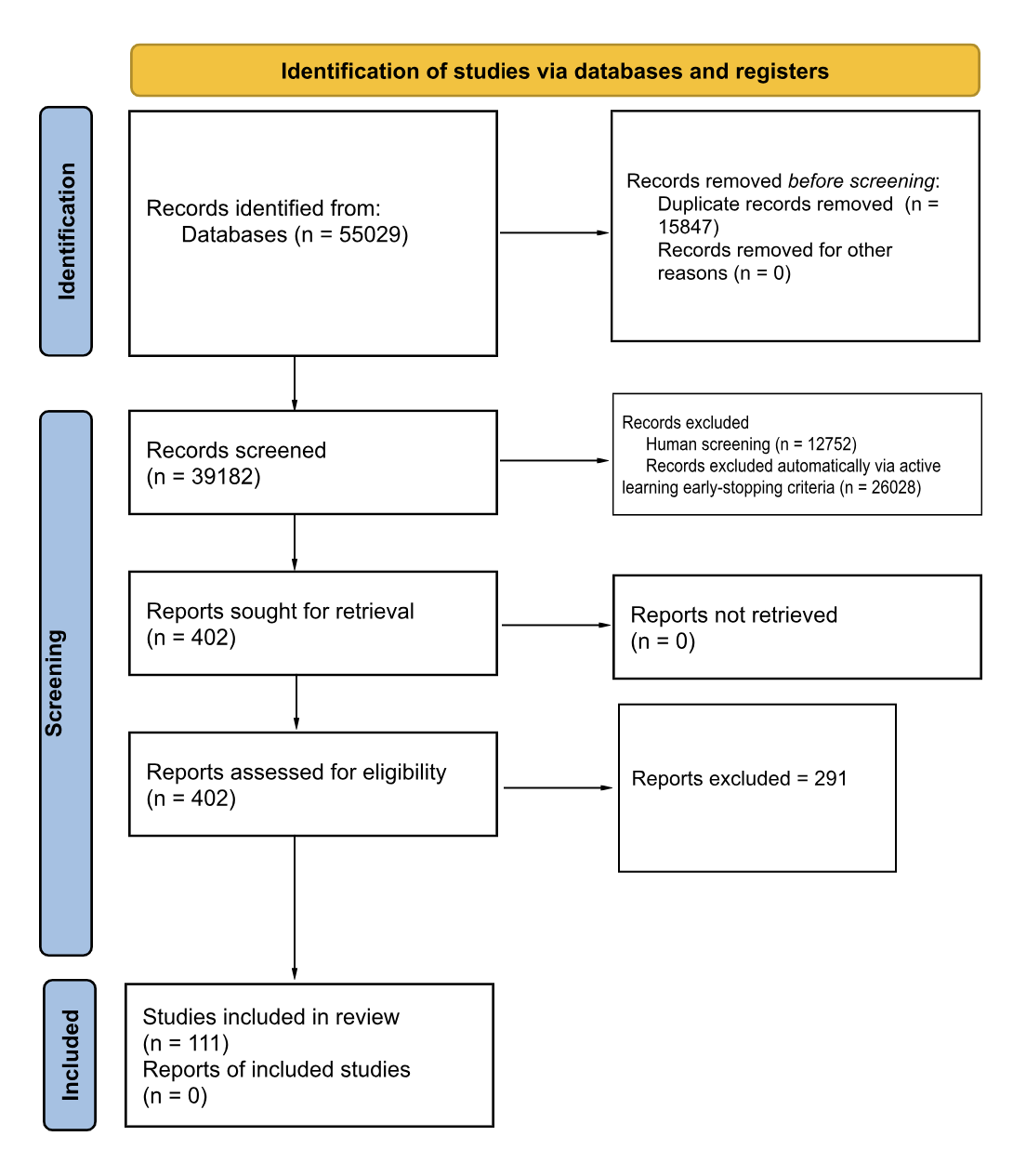

**Fig. S1. PRISMA flow diagram of study screening process.**

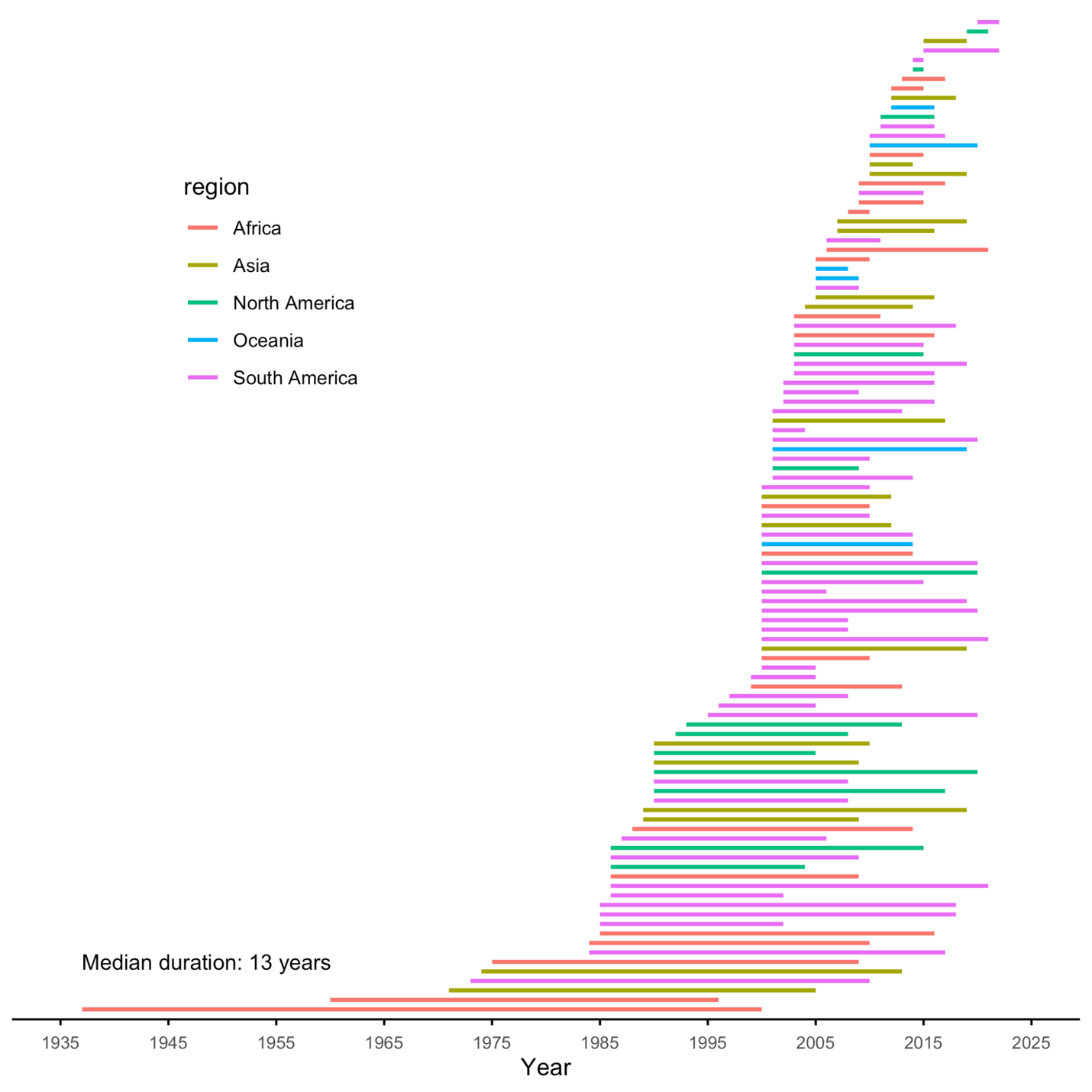

**Fig. S2.** Duration, start and end years of cases included in the meta-analytic comparisons of community-governed and conventionally governed areas (fully protected areas, partially protected areas, and areas outside protection).

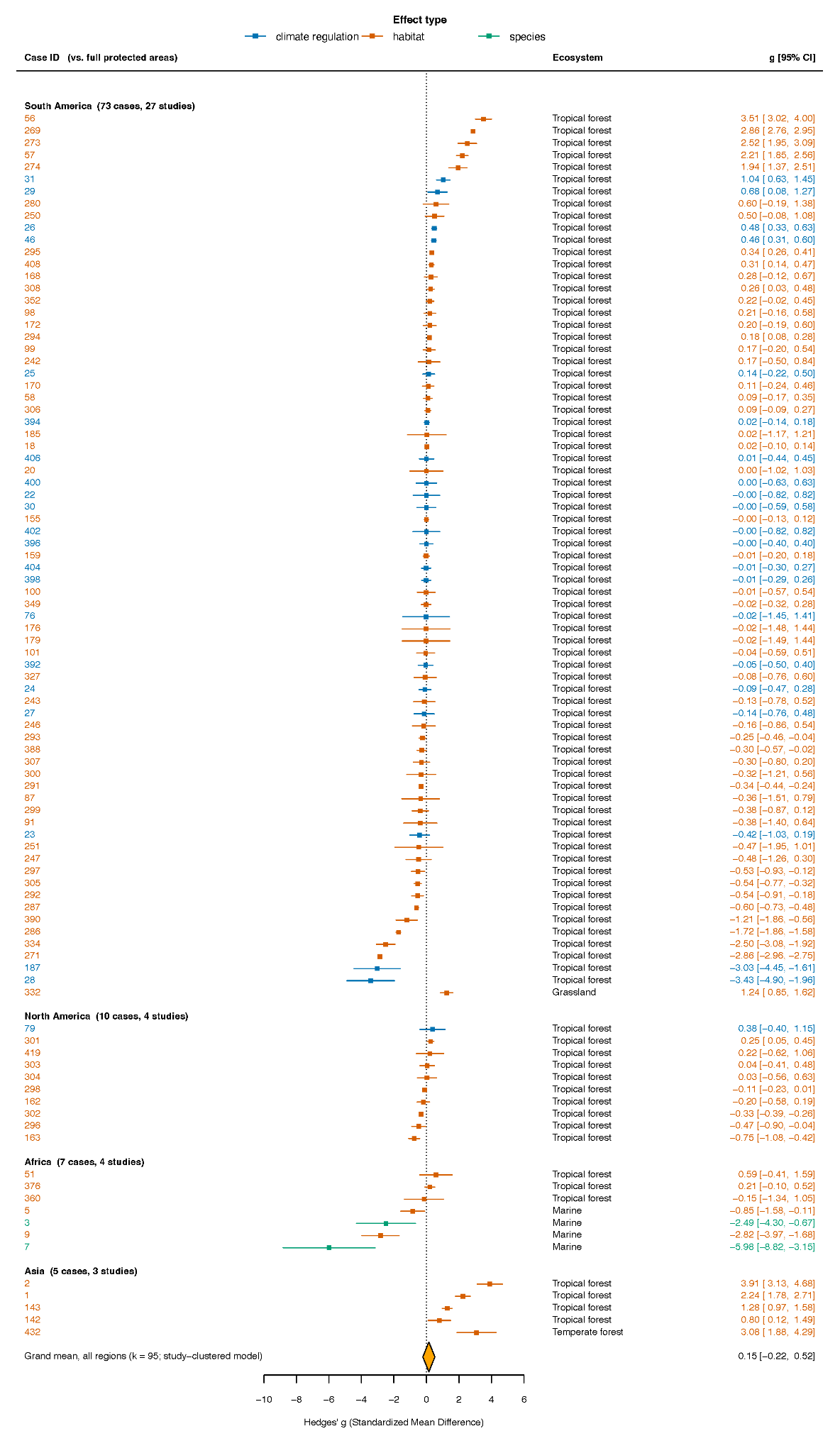

**Fig. S3.** Forest plot of all cases included in the meta-analyses of community governance vs. fully protected areas. Cases are grouped by region then ecosystem type, with colors of points (effect size, Hedges’ g) and whiskers (± 95% confidence intervals) denoting the type of ecological effect measured in each case. North America includes the countries of North and Central America. Orange diamond at the bottom of each figure shows the grand mean effect size computed in the main random-effects meta-analysis of cases clustered within studies, with 95% confidence interval denoted by the diamond width (as reported in Fig. 4).

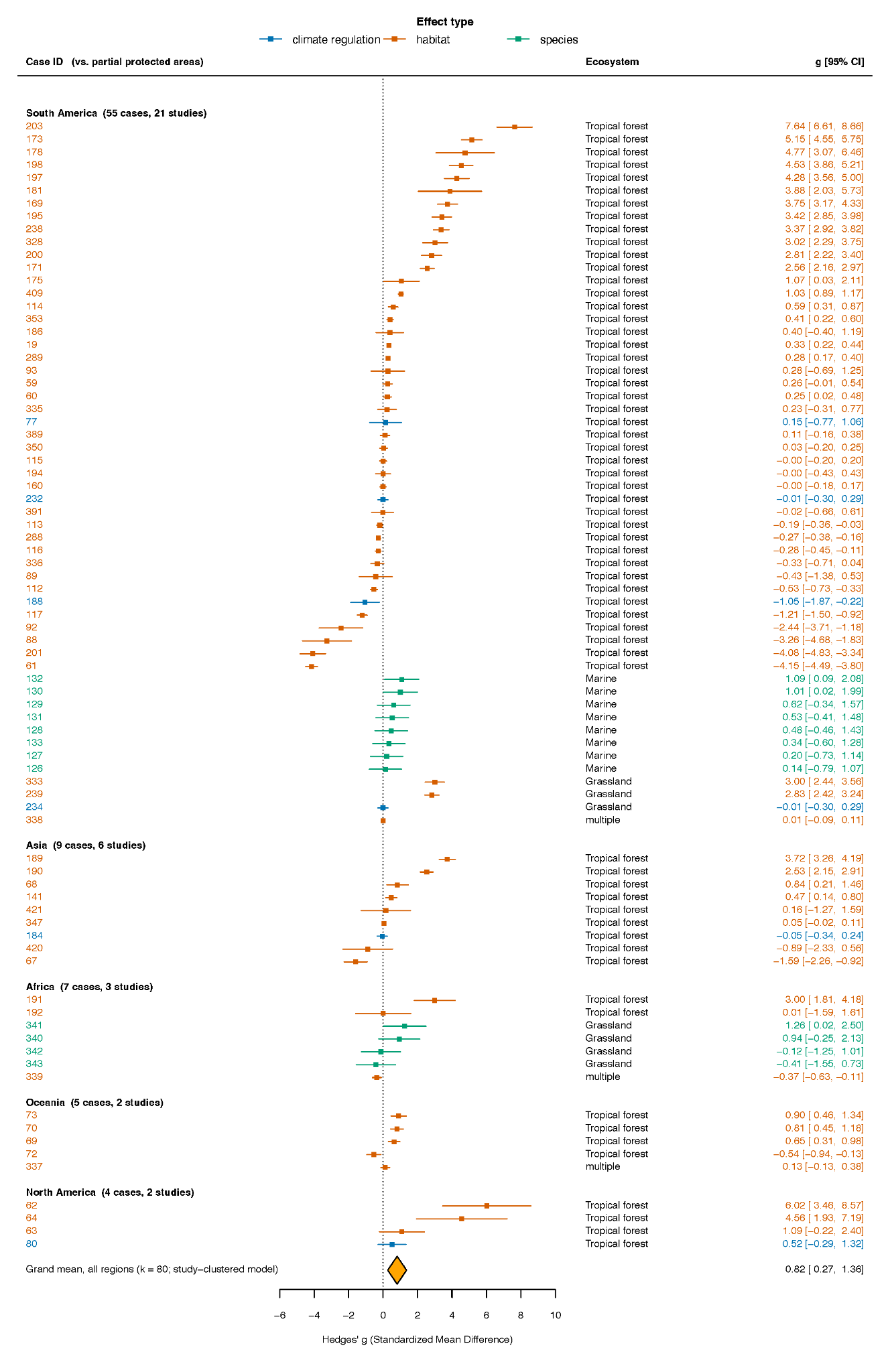

**Fig. S4.** Forest plot of all cases included in the meta-analyses of community governance vs. partially protected areas. Cases are grouped by region then ecosystem type, with colors of points (effect size, Hedges’ g) and whiskers (± 95% confidence intervals) denoting the type of ecological effect measured in each case. North America includes the countries of North and Central America. Orange diamond at the bottom of each figure shows the grand mean effect size computed in the main random-effects meta-analysis of cases clustered within studies, with 95% confidence interval denoted by the diamond width (as reported in Fig. 4).

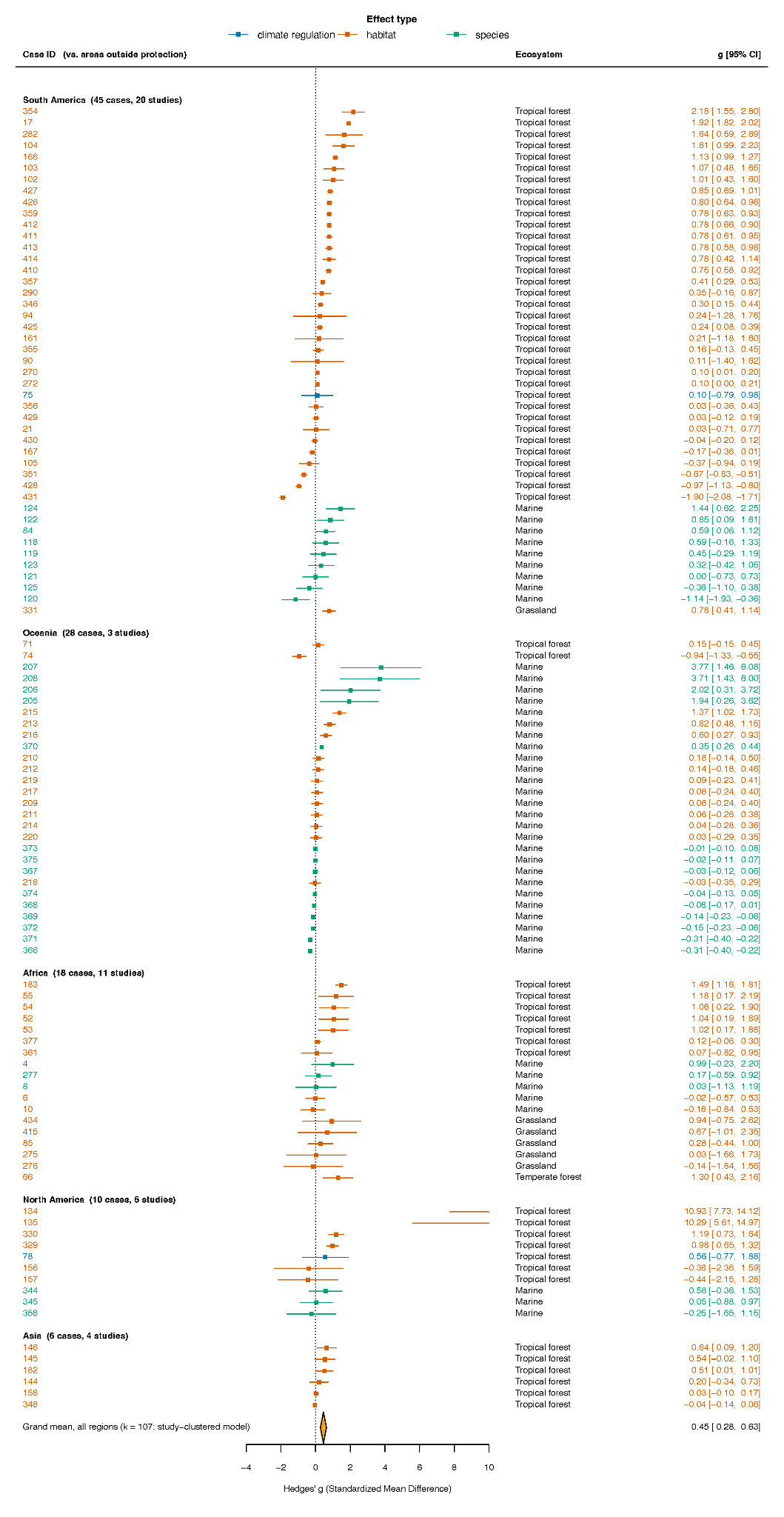

**Fig. S5.** Forest plot of all cases included in the meta-analyses of community governance vs. areas outside protection. Cases are grouped by region then ecosystem type, with colors of points (effect size, Hedges’ g) and whiskers (± 95% confidence intervals) denoting the type of ecological effect measured in each case. North America includes the countries of North and Central America. Orange diamond at the bottom of each figure shows the grand mean effect size computed in the main random-effects meta-analysis of cases clustered within studies, with 95% confidence interval denoted by the diamond width (as reported in Fig. 4).

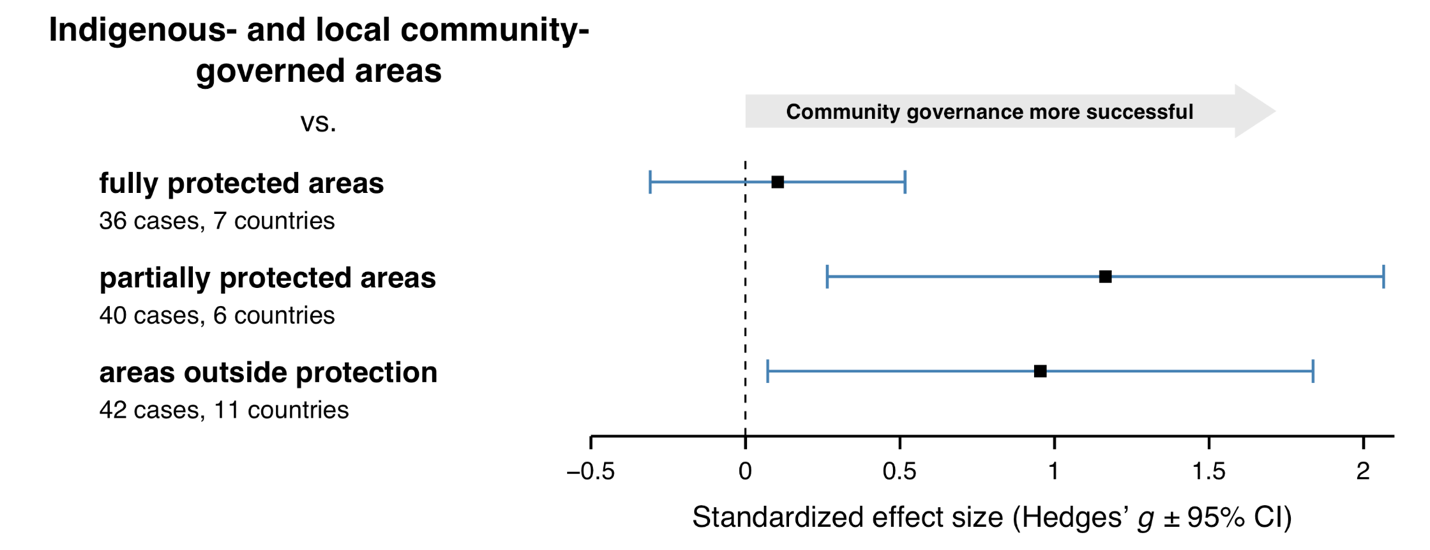

**Fig. S6.** **Effect sizes of community-governed areas from studies that employed matching**. Grand mean effect sizes and 95% confidence intervals for the subset of cases in each meta-analysis that employed matching techniques. From top to bottom, results of comparisons of ecological outcomes over time of community-governed areas to each comparison type (fully protected, partially protected areas, areas outside protection); and of titled community lands compared to untitled lands. Numbers of cases and countries in which cases were studied are reported for each comparison. Mean standardized effect sizes (Hedges’ g) are indicated by black squares, with 95% confidence intervals (CI) indicated by bar widths. Values to the right of zero indicate that community-governed area(s) had more positive ecological outcomes than comparison areas. The vertical dashed line indicates an effect size of zero. Effect sizes are significant where the confidence intervals do not overlap zero.

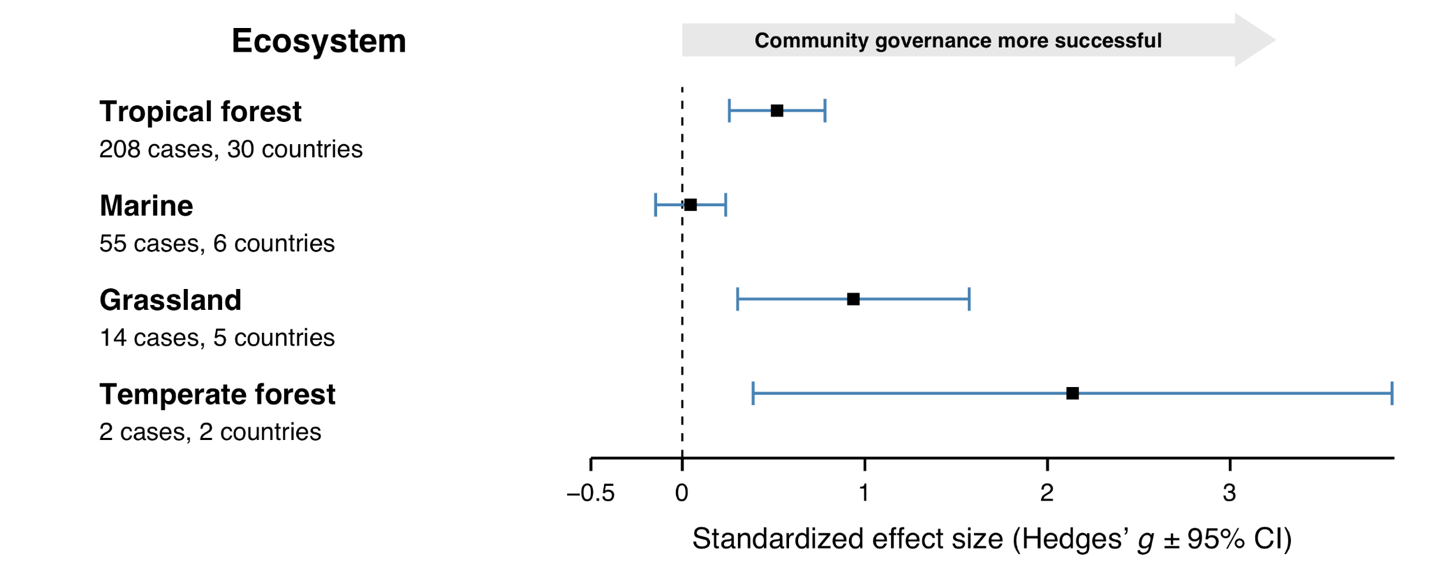

**Fig. S7. Effect sizes by ecosystem type of community-governed areas** against the pooled set of conventionally governed comparison types (fully protected areas, partially protected areas, and areas outside protection). Mean standardized effect sizes (Hedges’ g) are indicated by black squares, with 95% confidence intervals (CI) indicated by bar widths. Values to the right of zero indicate that community-governed area(s) had more positive ecological outcomes than comparison areas. The vertical dashed line indicates an effect size of zero. Effect sizes are significant where the confidence intervals do not overlap zero.

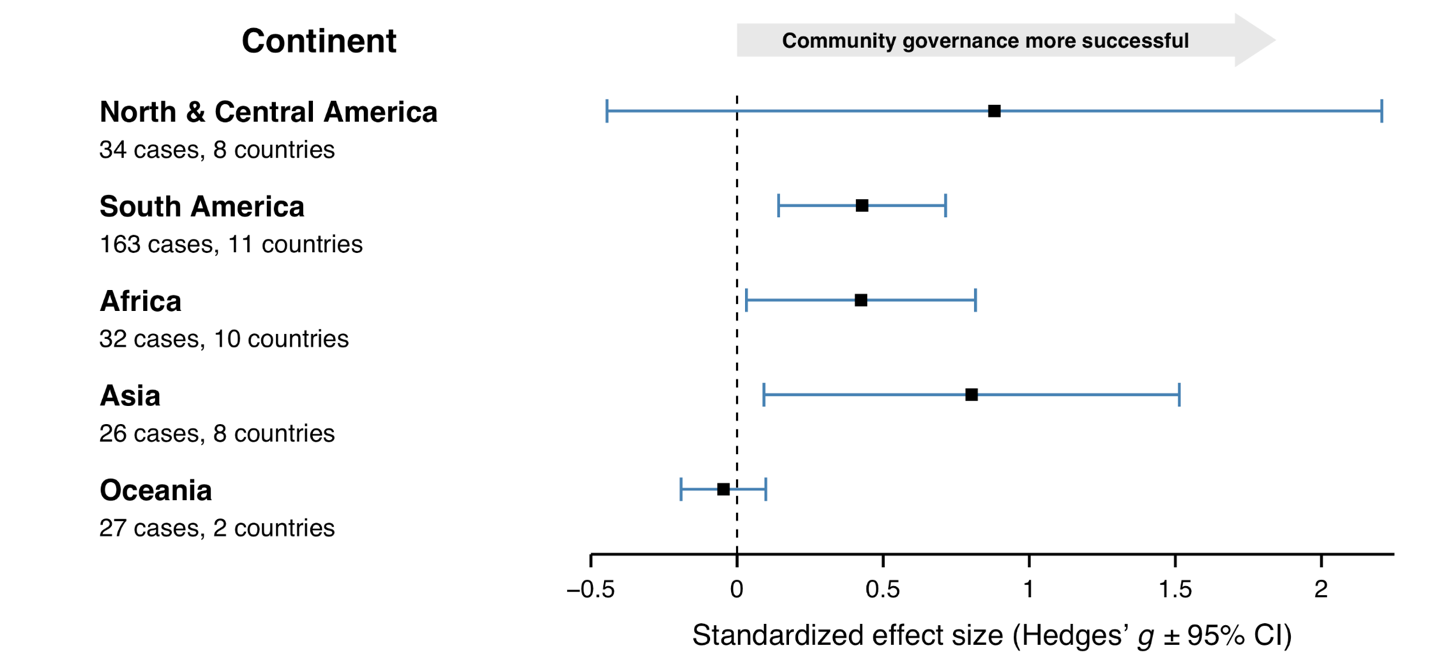

**Fig. S8. Effect sizes by region of community-governed areas** against the pooled set of conventionally governed comparison types (fully protected areas, partially protected areas, and areas outside protection). Mean standardized effect sizes (Hedges’ g) are indicated by black squares, with 95% confidence intervals (CI) indicated by bar widths. Values to the right of zero indicate that community-governed area(s) had more positive ecological outcomes than comparison areas. The vertical dashed line indicates an effect size of zero. Effect sizes are significant where the confidence intervals do not overlap zero.

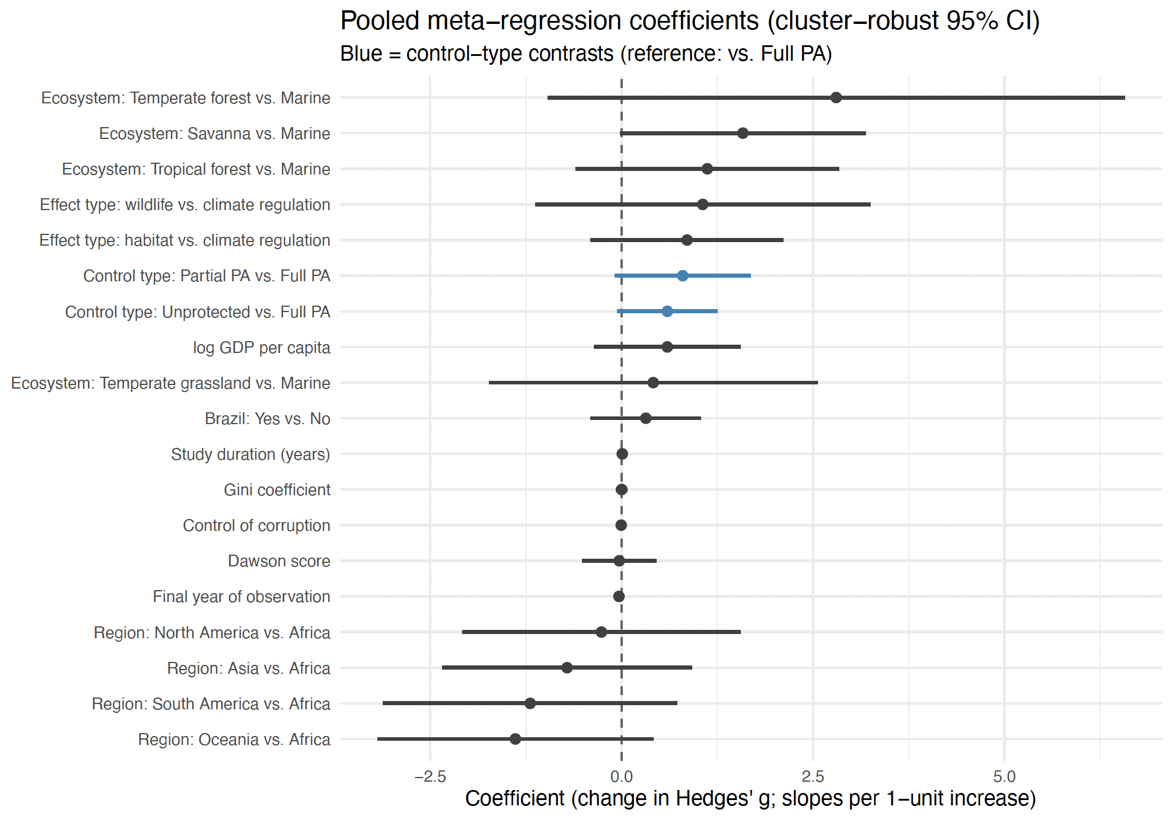

**Fig. S9: Meta-regression analysis coefficients** with 95% confidence intervals for pooled comparison of community-governed and conventionally-governed areas (fully protected areas, partially protected areas, and areas outside protection).

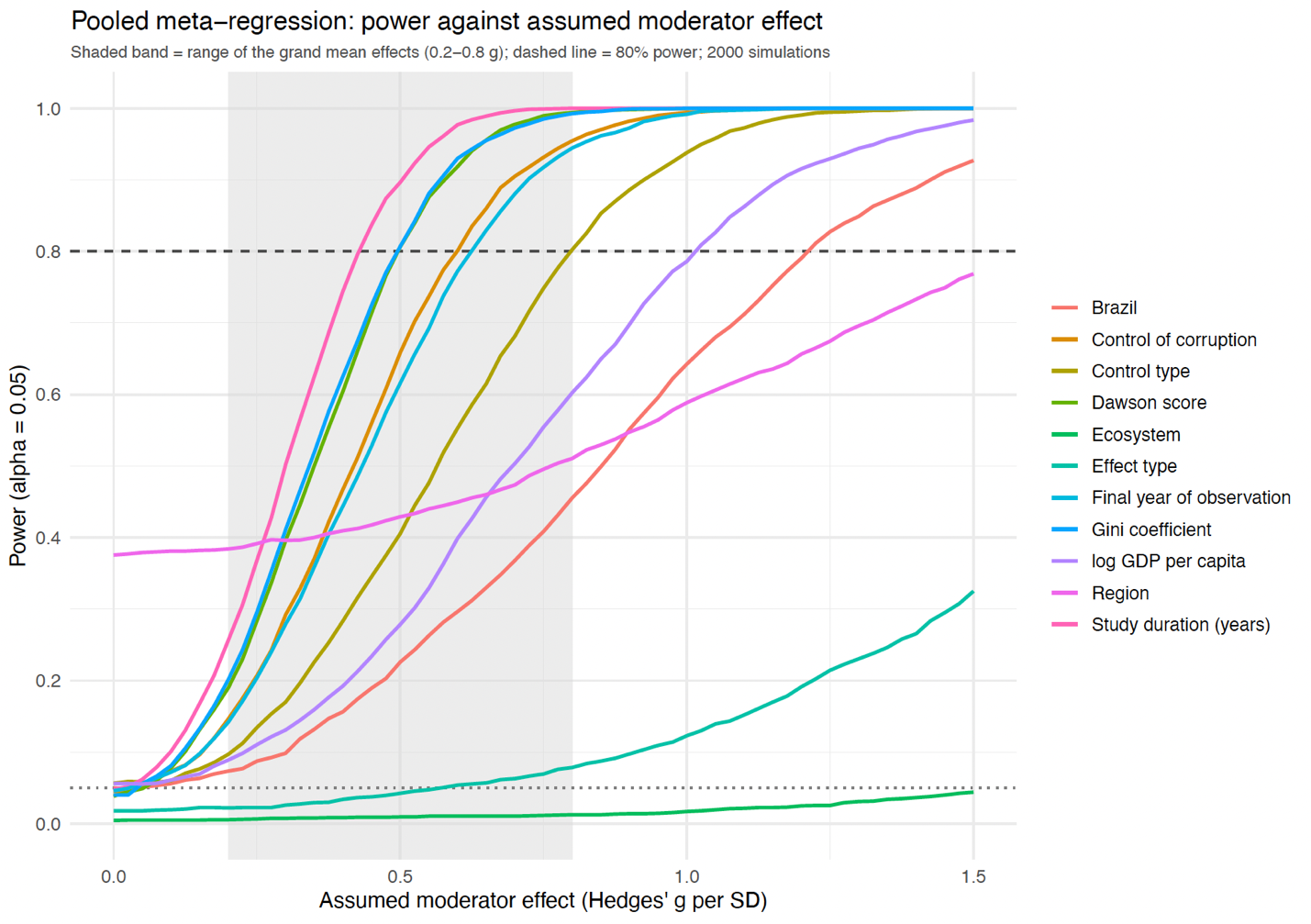

**Fig. S10. Simulated power curves for the meta-regression** pooled across the three comparison  types (fully protected areas, partially protected areas, and areas outside protection) for each moderator variable (n=2,000 runs). Dashed line at power=0.8 indicates at what level of influence on effect size (standardized difference between community-governed and conventionally-governed groups) the meta-regression had an 80% likelihood of detecting a significant influence. The gray band shows the range of overall grand mean effects (Hedges’ g) detected in the three meta-analyses comparing community-governed to conventionally-governed areas.

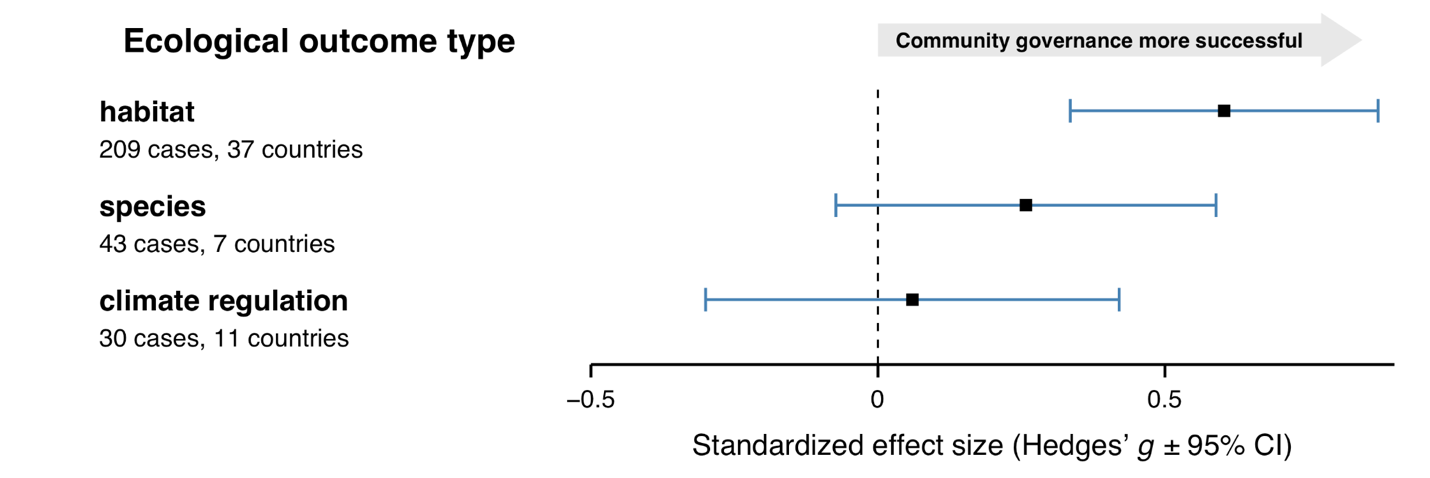

**Fig. S11. Effect sizes by ecological outcome variable type of Indigenous- and community-governed areas** against the pooled set of conventionally governed comparison types (fully protected areas, partially protected areas, and areas outside protection). Mean standardized effect sizes (Hedges’ g) are indicated by black squares, with 95% confidence intervals (CI) indicated by bar widths. Values to the right of zero indicate that Indigenous or community-governed area(s) had more positive ecological outcomes than comparison areas. The vertical dashed line indicates an effect size of zero. Effect sizes are significant where the confidence intervals do not overlap zero. Despite our categorization, we note that most habitat-scale variables (forest cover, vegetation greenness etc.) are expected to also contribute positively to climate regulation and to species outcomes.

**Supplementary Tables**

Table S1. Summary meta-analysis results for main, multi-level  random-effects models of effect size Hedges’ g, with cases nested within studies.

| Comparison | Case set | Mean standardized effect size (g) | Standard error | 95% CI interval | p-value |
| --- | --- | --- | --- | --- | --- |
| Community-governed vs. fully protected areas | All cases, n=95 | 0.150 | 0.189 | [-0.221,  0.521] | 0.429 |
|  | Matched cases, n=36 | 0.104 |  | [-0.308, 0.517] |  |
| Community-governed vs. partially protected areas | All cases, n=80 | 0.816 | 0.280 | [0.268,1.36] | 0.0035 |
|  | Matched cases, n=40 | 1.165 |  | [0.265, 2.07] |  |
| Community-governed vs. areas outside protection | All cases,  n=107 | 0.453 | 0.0894 | [0.278,0.628] | <0.0001 |
|  | Matched cases, n=42 | 0.954 |  | [0.072, 1.84] |  |
| Titled vs. untitled community-governed areas | All cases, n=31 | 0.437 | 0.136 | [0.171, 0.703] | 0.0013 |
|  | Matched cases, n=15 | 0.271 |  | [-0.131, 0.674] |  |

Table S2. Vote counts of ecological outcome over time by case in each meta-analysis (number of unique studies in parentheses).

| Test group | Comparison group | Test group did better | Comparison group did better | Tie between groups |
| --- | --- | --- | --- | --- |
| Community-governed areas (n=123) | Fully protected areas | 59 *(31)* | 64 *(28)* | 0 |
| Community-governed areas (n=98) | Partially protected areas | 63 *(31)* | 32 *(21)* | 3 *(2)* |
| Community-governed areas (n=187) | Areas outside protection | 133 *(59)* | 54 *(27)* | 0 |
| Community-governed areas with title (n=32) | Community-governed areas without title | 26 *(11)* | 6 *(5)* | 0 |

Table S3. Results of modified, multilevel Egger tests for publication bias; and trim-and-fill analysis for missing studies, with random-effects model refitted to include imputed missing cases. See Materials and Methods for details.

| Comparison | correlation of g with original SE, with modified SE | Multilevel PET slope (SE, p) | Multilevel PEESE slope (SE, p) | estimated missing studies (side, SE) | Refitted model with missing cases imputed: g (p, 95% CI) |
| --- | --- | --- | --- | --- | --- |
| Community-governed vs. fully protected areas | -0.40, -0.25 | -1.7 (0.99, p=0.11) | -2.7 (1.4, p=0.086) | 30 (right side, 6.22) | g=0.506 (p<0.0001, 0.259-0.753) |
| Community-governed vs. partially protected areas | 0.30, 0.047 | 0.33 (1.54, p=0.83) | 0.094 (2.09, p=0.97) | 0 | N/A |
| Community-governed vs. areas outside protection | 0.69, 0.22 | 2.9 (1.4, p=0.12) | 3.0 (1.8, p=0.19) | 15 (left side, 6.75) | g=0.218 (p=0.0089, 0.0547-0.381) |
| Titled vs. untitled community lands | 0.043, 0.014 | 0.042 (0.38, p=0.92) | 0.072 (0.30, p=0.85) | 0 | N/A |
